# Measurement of the torque in braided DNAs using a thermodynamic Maxwell relation

**DOI:** 10.1101/2020.02.07.938209

**Authors:** Botao Xiao, Sumitabha Brahmachari, Yang Liu, Ke Ding, John F. Marko

## Abstract

Braided DNAs are significant structural intermediates in cellular processes, yet little has been experimentally demonstrated about their higher-order structure and twisting torques. We use magnetic tweezers to measure braid extensions at forces ranging from 0.3 to 8 piconewtons, and then apply a thermodynamic Maxwell relation to calculate the torque. Experimentally inferred torques in unbuckled braids take on values up to 76 pN·nm, which depends on force, and inter-tether distance. As predicted using a statistical mechanical model, the twist modulus of the braids increases with catenation prior to buckling or formation of plectoneme, and is comparable to that of single DNA.

---

Millimeter-to centimeter-long double helical DNA molecules are known to form supercoils in the micron-sized cell [1,2]. The intertwining of two DNAs is frequently encountered in replication, recombination, and chromatid segregation [3–5]. An example of such intertwining occurs during replication where DNA strands are duplicated following the chiral double helix, resulting in rotational motions and torsional strains [6]: the two DNAs intertwine around each other to form a “braid” structure. We note that while not braids in the strict mathematical sense of the term, these 2-ply structures studied here have been conventionally called braids in the DNA micromanipulation literature [7].

Torque is a key physical parameter involved in DNA organization, affecting activities of enzymes such as topoisomerases, recombinases and polymerases [4,8–11], and serving as a key mechanical regulator in a multitude of cellular processes. For example, a resisting torque was shown to slow down and even stop transcription by RNA polymerase [12]. The torque in individual double-helix DNAs has been studied using single molecule techniques such as optical [13–15] and magnetic tweezers [16,17]. The twist modulus and phase transition-like behaviors of single DNAs have also been described by theoretical studies [18,19], in good accord with experiments. However, the torque in a double-DNA braid has never been measured experimentally.

The relation between extension and catenation of braided DNAs has been studied using magnetic tweezers (MT) under a few forces [3,20], though systematic studies over a broad range of forces have not been carried out. A model proposed by Kornyshev and Leikin suggests that left-handed DNA braids may undergo “collapse” and behave differently from right-handed DNA braids [21–23], and an electrically charged semi-flexible polymer model predicted a symmetry between positive and negative DNA braids of physiologically relevant braiding density [24]. However, the predictions of these models remain not fully experimentally explored. A further, biophysical question is that whether braided DNAs have a critical buckling condition, which may relate to the phase coexistence phenomena observed for twisting and stretching a single DNA helix [18]. However, it is challenging to measure large numbers of DNA braids in a single experimental flow cell, as the numbers of DNA braids in a given experiment is typically much fewer than that of single DNAs [25].

It is known that structures of intertwined DNA molecules affect activities and functions of braid-interacting proteins. Efforts have been made to study DNA coil-coil and RNA coil-coil structures by X-ray crystallography, but only tens of base pairs could be resolved [26]. Previous studies have indicated that at high catenation numbers, twisted DNA braids undergo a mechanical instability, likely resulting in the formation of plectonemic-braided structures. Theoretical work has predicted the critical braid torque [23] at this instability, but this has not yet been measured.

In this letter, the relationship between the extension and catenation density was systematically measured with holding forces ranging from 0.3 to 8 pN, which are within the physiological range. We used 5.9 kb DNA fragments derived from mM502, 48.5 kb lambda phage DNA, and 6 kb linear fragments derived from plasmids pFOS1, via PCR reactions using biotin-and digoxigenin-labeled primers [3–4]. The labeled DNA strands were incubated and bound with one end to streptavidin-coated paramagnetic beads of which the diameter was 1 or 2.8 μm (Dynabeads), and with the other end to an anti-digoxigenin coated glass surface. All experiments were performed in the phosphate-buffered saline (PBS) containing 150 mM NaCl and 0.5 mg/ml bovine serum albumin (BSA), and at room temperature (~25 °C). We put this DNA sample onto the MT to stretch the microbeads and rotated them to braid the two DNA molecules as previously described [3].

Tethers in the flow cell were tested for appropriate force-extension behaviors and selected. Braid extension measured under zero catenation at various forces displayed wormlike chain behavior with a persistence length of *A_eff_* ≈ 25 nm, or about half that of the *A* = 50 nm characteristic of a single DNA molecule. This follows from the fact that the braids at zero catenation are two parallel DNA molecules, and when stretched one sees twice the usual spring constant [7, 29]. We also used the condition wherein the spring constant was double that of a single DNA to distinguish a braid among the experimental samples which contain multiple DNA tethers that would feature effective persistence length less than a half of a single DNA.

At each stretching force, the two DNA molecules were braided around each other by rotating the magnets in a clock-wise (Figure 1b-d) or counter-clockwise direction. By changing the Ca from 0 to ±1/2, the extension *z* exhibited a sharp drop. When the Ca was above 1/2, the extension decreased gradually when the molecules were further catenated (Figure 1g). The extension data points at each force were fit by a 2^nd^ order polynomial function: *z* = *x*_1_*Ca*^2^ + *x*_2_*Ca* + *x*_3_.

**Figure 1.**
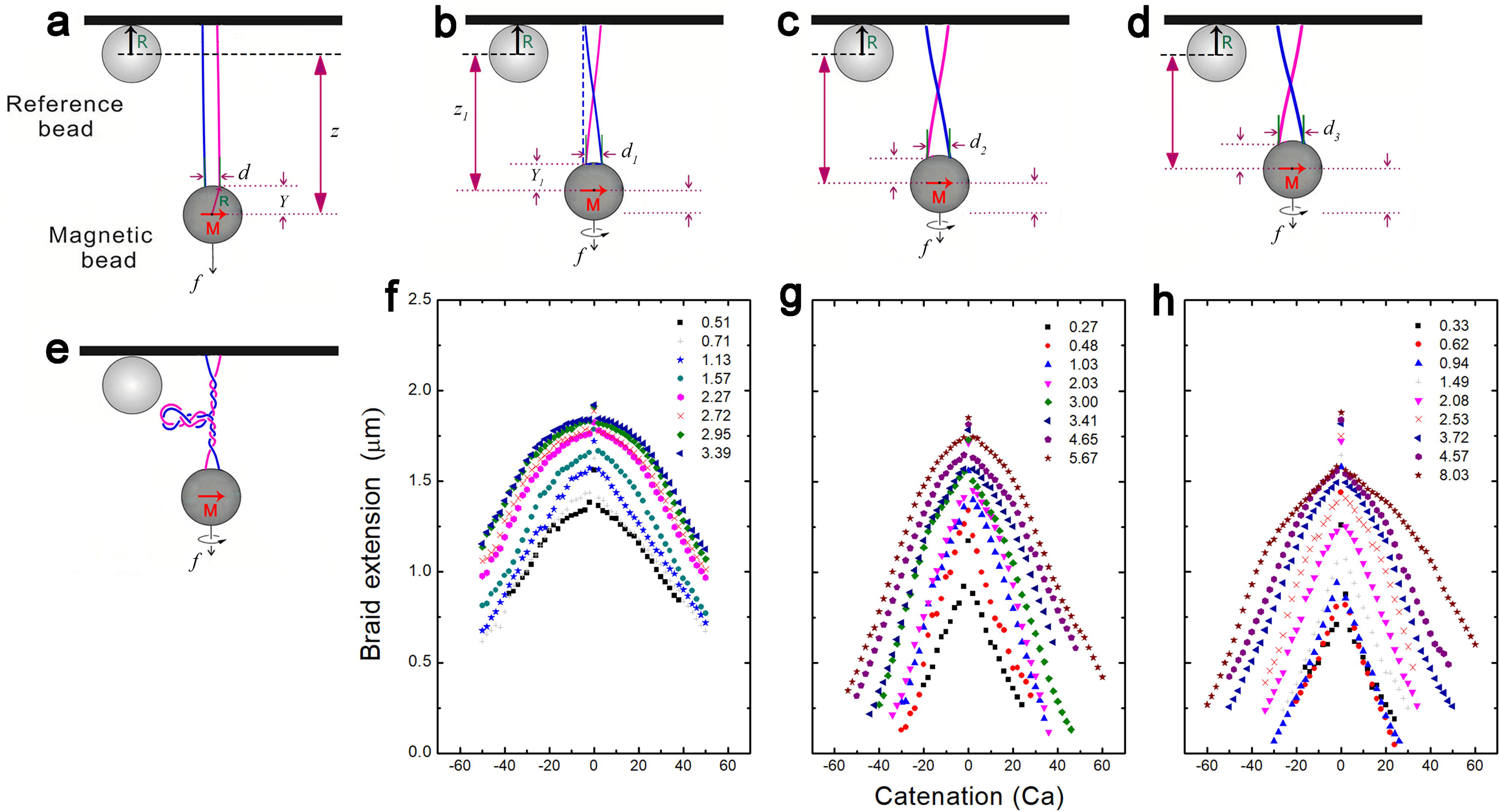
Braiding of two DNA molecules. **a** A schematic shows that two DNA molecules are anchored to a magnetic bead, and they can be braided around each other by rotating the bead. *R*, *z* and *d* indicate the radius of the magnetic bead, extension, and intertether distance of the DNA braid, respectively. **b-d** Schematics of the small (**b**), mid (**c**) and large (**d**) intertether distance braids show that there is a sharp decrease in extension for the ±1/2 turn, followed by gentle decreases when the turn is larger than 1/2 or smaller than −1/2. **e** A plectoneme forms as the Ca further increases. **f-g** For the small (*d=*0.26 *L*) (**f**), mid (0.31 *L*) (**g**) and large (0.54 *L*) (**h**) intertether distance braids, the extensions under positive and negative Ca are symmetric about Ca=0 which is the buckling transition point. Each data point was obtained by averaging ~100 times of fast extension measurements.

At higher values of catenation, the extension decreased almost linearly and were off the polynomial line (Figure 2a). Different DNA extensions were normalized to a relative extension and catenation was normalized to the catenation density (*σ*) using the total DNA linking number: *σ* =*Ca h* /*L*, where h=3.6 nm is the DNA helix repeat and *L* is the total DNA contour length. For example, the value of *σ* at Ca=56 was 0.997 for the mM502 fragment of which the contour length (*L*) was 1.97 μm. The force-extension relations for positive and negative Ca were found to be symmetric about Ca=0 (Figure 1f-h). The catenation-extension curves of different pairs of braids at the same force did not overlap (Figure 1f-h), which was due to their different intertether distances (*d*). The extension shortening between Ca=0 and Ca=1/2 was used to calculate the intertether distance (*d*) by the Pythagorean theorem [30]. The *d* values were found to be 0.52, 0.62, and 1.06 μm in Figure 1f-h, respectively. The *d/L* ratios were 0.26, 0.31, and 0.54 for the mM502 fragment. At the same force, the extensions of a narrow intertether (*d/L*=0.26) braid were slightly longer than those of the wider intertether (*d/L*=0.31) braid or the widest intertether (*d/L*=0.54) braid (Figure 1f-h).

We calculated the change of torque using a thermodynamic Maxwell relation. For braids in the buffer, the differential of the thermodynamic potential *E* can be described in terms of the Gibbs isotherm [27]:

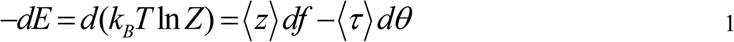

**Figure 2.**
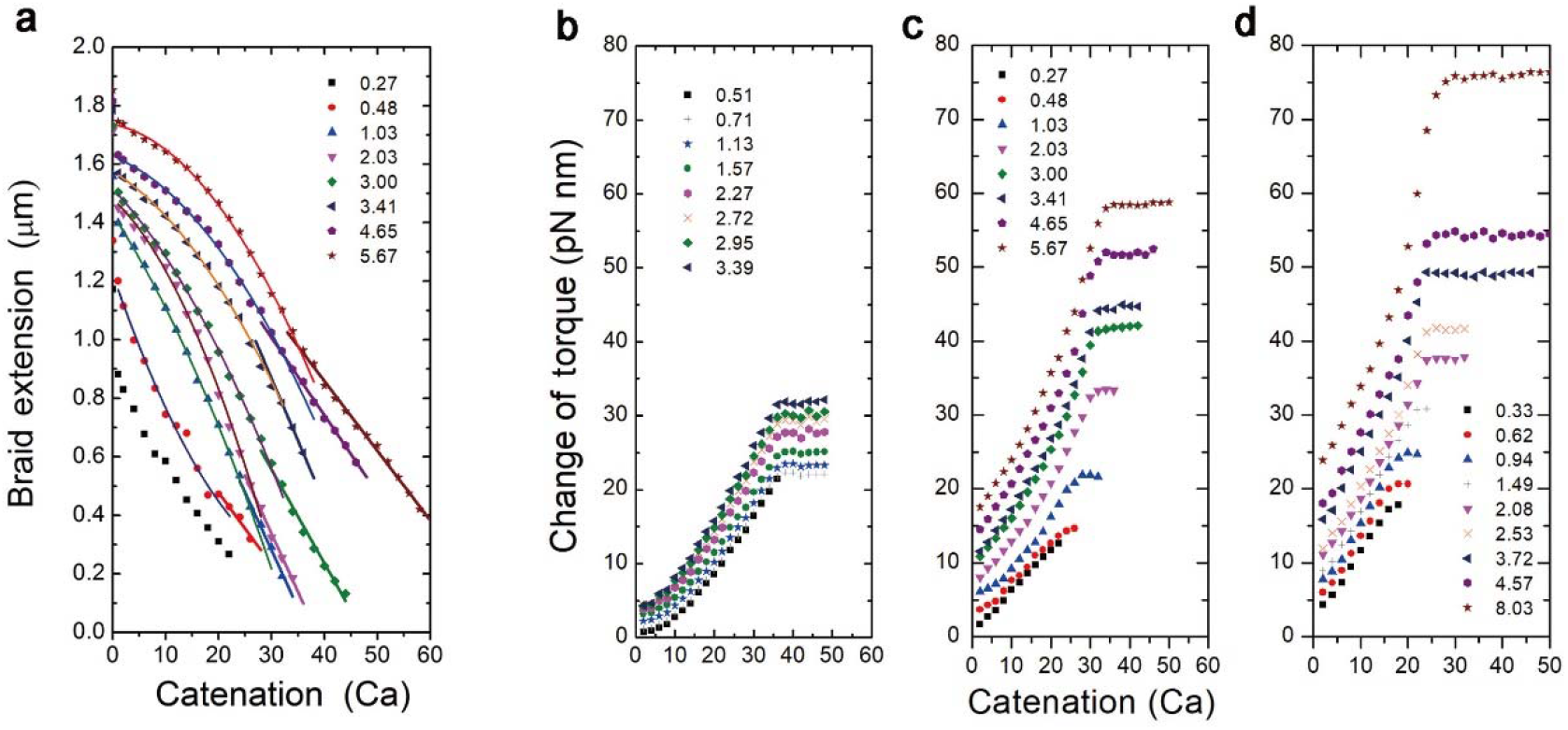
**a** The extension-catenation curves of the mid intertether distance (0.31 *L*) braid under different forces (Figure 1g) are fitted by a polynomial line, and the tails are fitted by a straight line. Due to the symmetry of the extensions relative to negative and positive Ca, the negative Ca data points are not shown here. **b–d** For the small (*d=*0.26 *L*) (**b**), mid (0.31 *L*) (**c**) and large (0.54 *L*) (**d**) intertether distance braids, change of torque τ increases with the catenation until reaches a plateau.

For the *f-z* measurements at fixed *θ* in an equilibrium state, the change in torque τ can be obtained by integration as the force increases from *f*_0_ to *f* [16,27]:

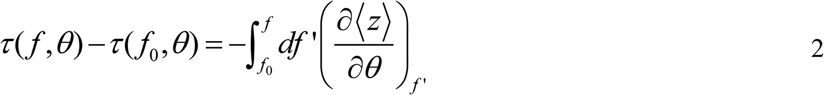

Here *df* is the difference between adjacent forces, the rotational angle *θ*=*2πCa*.

For the wider intertether braid (*d/L*=0.31) held by a 0.27 pN force, the change of torque was 1.7 pN·nm when the two DNAs were catenated by 1 turn (Figure 2c), and the change of torque increased to 13 pN·nm when the catenation was further increased to 22. At a stronger holding force of 5.67 pN, the change of torque was found to be 18 pN·nm when the Ca increased from 0 to 22. The torque increased with Ca until reached a plateau of ~58 pN·nm at Ca=52.

For the widest intertether braid (*d/L*=0.54) held by an 8.03 pN force, the torque increased from 0 to a peak of 76 pN·nm when the catenation was increased from 0 to Ca=30 or *σ*=0.53 (Figure 2d). At 2 pN force and Ca=20, the torques were 31, 21 and 15 pN·nm in braids of large (0.54 *L*), mid (0.31 *L*) and small (0.26 *L*) intertether distances, respectively (Figure 2b).

Theoretical calculations generating the extension-catenation curves (Figures 3d and 4d) were carried out based on a statistical-mechanical model for braids (See Supplementary Material) [24,31]. The torques from the model (Figures 3e and 4e) were obtained from the derivative of the free energy as a function of catenation. Torques from the experiments under the same conditions as the model are shown in Figures 3b and 4b. In general, the plots from the model agree with the experimental results calculated by the Maxwell relation, including the increasing torsional modulus of braids with catenation before the buckling point (Figures 3 and 4).

**Figure 3.**
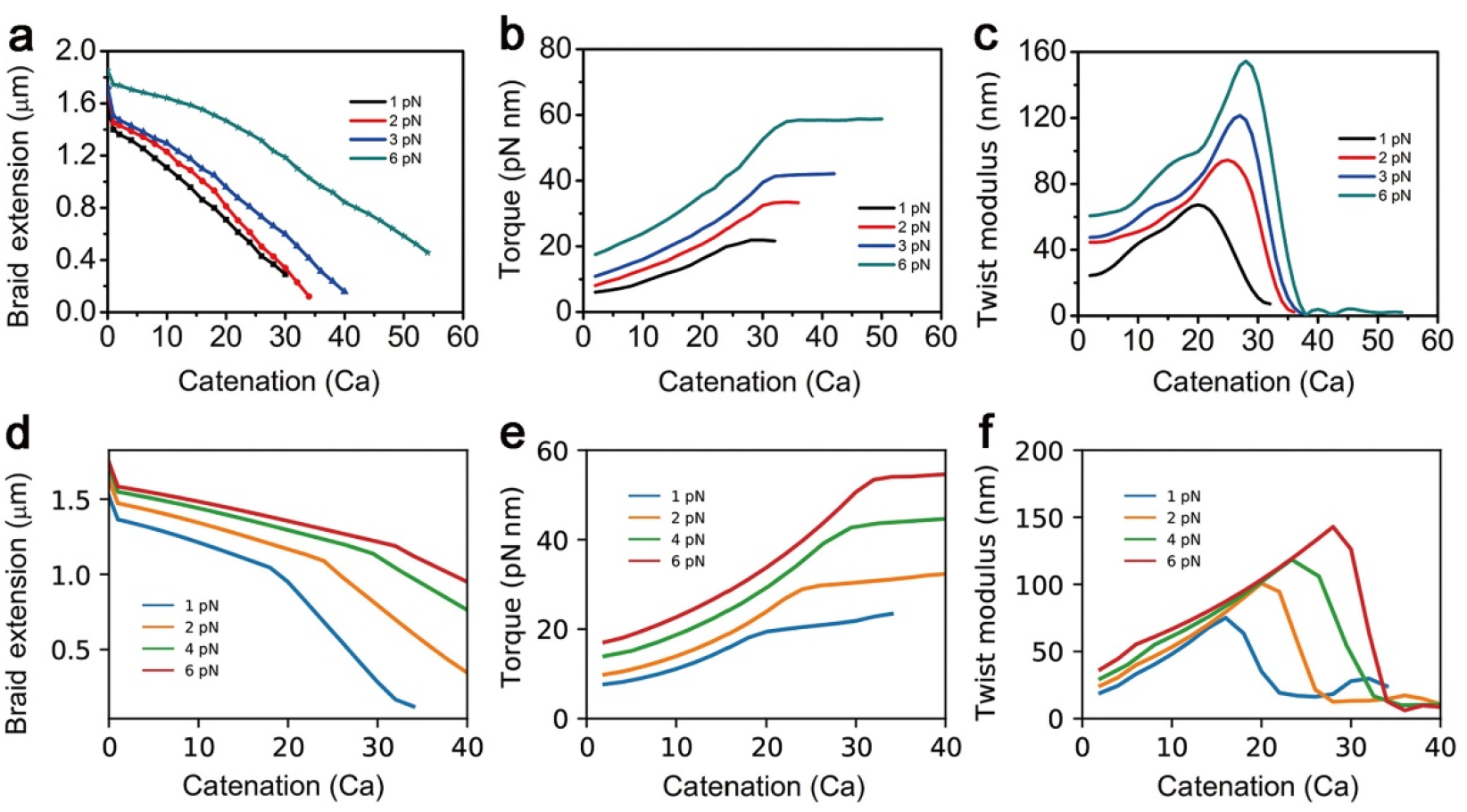
**a-c** Experimental extension (**a**), torque (**b**) and twist modulus (**c**) of a braid under different catenations with net stretching forces of 1, 2, 3 and 6 pN. The DNA is 5.9 kb. **d-f** The model calculation results of the extension (**d**), torque (**e**) and twist modulus (**f**) under different catenations with holding forces of 1, 2, 4 and 6 pN. The trends in (**a**), (**b**) and (**c**) are similar to those in (**d**), (**e**) and (**f**), respectively.

**Figure 4.**
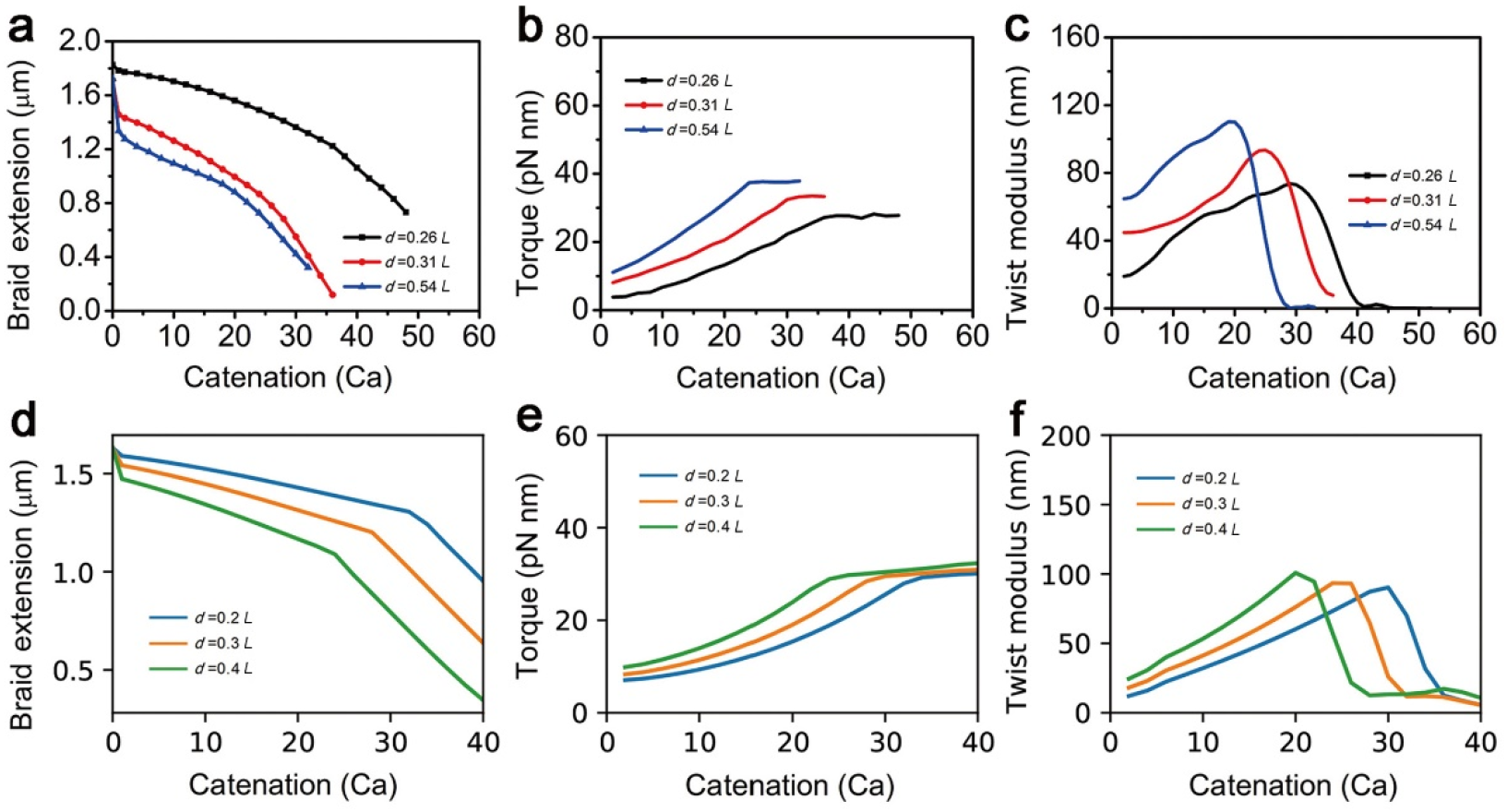
**a-c** Experimental extension (**a**), torque (**b**), and twist modulus (**c**) of small (*d=*0.26 *L*), mid (0.31 *L*) and large (0.54 *L*) intertether distance braids under different catenations with a net stretching force of 2 pN applied on the braids. The DNA is 5.9 kb. **d-f** The model calculation results of the extension (**d**), torque (**e**) and twist modulus (**f**) of small (*d=*0.2 *L*), mid (0.3 *L*) and large (0.4 *L*) intertether distance braids under different catenations with a holding force of 2 pN. The DNA is 5.7 kb. The trends in (**a**), (**b**) and (**c**) are similar to those in (**d**), (**e**) and (**f**), respectively.

The twist or torsional modulus *C* was calculated using the equation 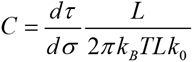, where *K_B_* is the Boltzmann constant, *T* is the absolute temperature, and *Lk*_0_ is the linking number of the DNA [15,16]. The modulus from the experimental data increased with the catenation until it reached a peak (Figures 3c and 4c). For the three braids of different intertether distances (Figure 2b-d), the peak values of *C* at 2 pN were 73.6, 93.4, and 110.2 nm, respectively (Figure 4c). For the same braid in Figure 2b but at different forces, the peak values of *C* were found to be 67.2 nm at 1 pN, 94.5 nm at 2 pN, 121.4 nm at 3 pN, and 154.5 nm at 6 pN (Figure 3c).

The theoretical twist modulus of DNA braids was also obtained from the derivative of the torque as a function of catenation (Figures 3e and 4e). The twist modulus increased with catenation and exhibited a “twist-stiffening” behavior [24], suggesting that the braids with a higher linear density of catenation had a stiffer twist response. The modulus is expected to scale approximately as the quadratic power of catenation (~Ca^2^), which is derived from a cubic torque and a quartic free energy. Note that the modulus is symmetric with respect to a sign change in Ca, suggesting similar stiffness in positively and negatively catenated braids. This behavior arises from the symmetry in the free energy of braid with respect to positive and negative catenations that generates extension curves which also have this mirror-symmetry with respect to Ca. The twist modulus decreased after the braid buckled (Figures 3e and 4e), suggesting the loss of twist rigidity in a buckled braid due to the absorption of excessive catenation via writhing of the braid into buckled plectoneme domains.

Twist stiffening response was stronger for braids of larger intertether distances than that for braids of small and mid intertether distances (Figure 4e). Braids with a larger intertether distance had larger end regions and consequently a smaller helically wrapped region. As a result, for a given catenation, the braids with a larger intertether distance would possess a higher linear density of catenation, leading to a steeper increase in torque with catenation.

The torque acting between two catenated DNAs was measured. Our results show that the torque increased with the catenation until reaching a plateau at the buckling point. The buckling torques of the braids were from 20 to 58 pN·nm, whereas the buckling torques of the single DNAs subject to twisting were up to 30 pN·nm [15,16]: the buckling torques for braids are comparable to, but somewhat larger than, single DNAs. The twist modulus of single DNA molecules is nearly constant with catenation at constant force [15,16]. By contrast, for DNA braids, we find that braid twist modulus increases with catenation before reaching the buckling point: this effect is theoretically expected [24] due to the gradual tightening of the braid, which generates an approximately Ca^4^ free energy density increase (plus lower-Ca-power corrections) due to confinement of wormlike chain bending fluctuations. A quantitatively similar behavior is seen in the experimental results, in the nonlinear increase in torque and corresponding increase in torsional modulus with catenation (Figures 3c and 4c). The modulus then decreases after buckling, theoretically expected due to the emergence of buckled domains that can absorb catenation in form of plectoneme writhe [24].

Overall, our experimental results agree well with theoretical calculation using the polymer model, although the experimental buckling points for braids of mid and large intertether distances (Figure 4b) slightly deviated from those predicted by theoretical modeling based on established models of DNA polymer statistical mechanics (Figure 4e). Recent work using an oscillating MT method estimated braid torques to be ~500 pN·nm, much larger than the torques reported here. This difference may be due to the large intertether distances, or the intrinsically non-equilibrium nature of the oscillating MT approach [32].

The rich dynamics of supercoiling of a single DNA molecule has been observed using single molecule fluorescent techniques [33], which observed highly dynamic and stochastic motions of small, supercoiled domains. Direct experimental investigation of braid dynamics is lacking, however, simulations studying plectoneme braids have observed dynamic features such as diffusion and hopping of plectoneme domains [34]. It has been observed that mechanical properties of the braid depend on temperature and salt condition [35]. On theoretical grounds it has been predicted that highly twisted braids should tend to form smaller plectonemic domains than supercoiled single DNAs due to the greater bulk (cross-sectional area) of braided DNAs [24]. Our experimental reports of the buckling dynamics of braid shave indirectly observed the effects of multiple small domains [31]. Future experiments using fluorescence visualization of braid supercoiling would be desirable to allow direct observation of the size and dynamics of braid-plectoneme domains.

The dependence of braid properties on physiochemical factors, along with the general differences between the braids and single DNAs, may influence cellular functions or the interactions between proteins and DNAs during replication, repair, and recombination, processes where interwound double helicies of the sort studied here can be expected to arise. A key question is how DNA with bound proteins, as will be found *in vivo*, will behave during interwinding, and to quantitatively establish how torque built up in DNA braids affects interwinding-acting enzymes including DNA-bending proteins [28,35], polymerases, topoisomerases [36–38], recombinases [3,4], and SMC complexes [39].

We thank Prof. Reid C. Johnson for providing the mM502 plasmid, and we thank Profs. Pei Guo, Jie Yan (National University of Singapore), Mian Long (Chinese Academy of Sciences), Yi Xiao and Jianhua Wu for helpful discussions. We thank Huiling Bai, Zhen Li, Dr. Hua Bai, Zhengming Wang, Xianbin Lei, and Qingshan Fu for technical assistance.

Research at SCUT and HUST was supported by the National Natural Science Foundation of China (11772133, 11372116) and the Fundamental Research Funds for the Central Universities (HUST 0118012051). Work at NU was supported by the NIH through grants R01-GM105847 and U54-CA193419.

## References

[1] V. Wood et al., Nature, 415, 871 (2002).

[2] J. Yan, R. Kawamura, and J.F. Marko, Phys. Rev. E, 71, 061905 (2005).

[3] B. Xiao, M.M. McLean, X. Lei, J.F. Marko, and R.C. Johnson, Sci. Rep., 6, 23697 (2016).

[4] H. Bai, M.X. Sun, P. Ghosh, G.F. Hatfull, N.D.F. Grindley, and J.F. Marko, Proc. Natl. Acad. Sci. U.S.A., 108, 7419 (2011).

[5] P. Cejka, J.L. Plank, C.C. Dombrowski, and S.C. Kowalczykowski, Mol. Cell, 47, 886 (2012).

[6] J.D. Moroz and P. Nelson, Proc. Natl. Acad. Sci. U.S.A., 94, 14418 (1997).

[7] T.R. Strick, J.F. Allemand, D. Bensimon, and V. Croquette, Biophys. J., 74, 2016 (1998).

[8] T.R. Strick, V. Croquette, and D. Bensimon, Nature, 404, 901 (2000).

[9] K.C. Neuman, G. Charvin, D. Bensimon, and V. Croquette, Proc. Natl. Acad. Sci. U.S.A., 106, 6986 (2009).

[10] S.C. Dillon, and C.J. Dorman, Nat. Rev. Microbiol., 8, 185 (2010).

[11] Y. Seol, A.H. Hardin, M.P. Strub, G. Charvin, and K.C. Neuman, Nucleic Acids Res., 41, 4640 (2013).

[12] J. Ma, L. Bai, and M.D. Wang, Science, 340, 1580 (2013).

[13] Z. Bryant, M.D. Stone, J. Gore, S.B. Smith, N.R. Cozzarelli, and C. Bustamante, Nature, 424, 338 (2003).

[14] S. Forth, C. Deufel, M.Y. Sheinin, B. Daniels, J.P. Sethna, and M.D. Wang, Phys. Rev. Lett., 100, 148301 (2008).

[15] M.Y. Sheinin, S. Forth, J.F. Marko, and M.D. Wang, Phys. Rev. Lett., 107, 108102 (2011).

[16] F. Mosconi, J.F. Allemand, D. Bensimon, and V. Croquette, Phys. Rev. Lett., 102, 078301 (2009).

[17] J. Lipfert, M.M. van Oene, M. Lee, F. Pedaci, and N.H. Dekker, Chem. Rev., 115, 1449 (2015).

[18] J.F. Marko and S. Neukirch, Phys. Rev. E, 88, 062722 (2013).

[19] J.F. Marko, Phys. Rev. E, 76, 021926 (2007).

[20] G. Charvin, A. Vologodskii, D. Bensimon, and V. Croquette, Biophys. J., 88, 4124 (2005).

[21] R. Cortini, A.A. Kornyshev, D.J. Lee, and S. Leikin, Biophys. J., 101, 875 (2011).

[22] D.J. Lee, J. Phys-Condens. Mat., 27, 145101 (2015).

[23] D.J. Lee, R. Cortini, A.P. Korte, E.L. Starostin, G.H.M. van der Heijden, and A.A. Kornyshev, Soft Matter, 9, 9833 (2013).

[24] S. Brahmachari and J.F. Marko, Phys. Rev. E, 95, 052401 (2017).

[25] I. De Vlaminck et al, Nano Lett., 11, 5489 (2011).

[26] J.R. Stagno, B.Y. Ma, J. Li, A.S. Altieri, R.A. Byrd, and X.H. Ji, Nat. Commun., 3, 901 (2012).

[27] H. Zhang and J.F. Marko, Phys. Rev. E, 77, 031916 (2008).

[28] B. Xiao, H. Zhang, R.C. Johnson, and J.F. Marko, Nucleic Acids Res., 39, 5568 (2011).

[29] J.F. Marko, Phys. Rev. E 55 1758 (1997).

[30] G. Charvin, D. Bensimon, and V. Croquette, Proc. Natl. Acad. Sci. U.S.A., 100, 9820 (2003).

[31] S. Brahmachari, K.H. Gunn, R.D. Giuntoli, A. Mondragon, and J.F. Marko, Phys. Rev. Lett., 119, 188103 (2017)

[32] C.J. Martinez-Santiago and E. Quinones, Chem. Phys., 502, 6 (2018).

[33] M.T.J. van Loenhout, M.V. de Grunt, and C. Dekker, Science, 338, 94 (2012).

[34] G. Forte, M. Caraglio, D. Marenduzzo, and E. Orlandini, Phys. Rev. E 99, 052503 (2019).

[35] B. Xiao, R.C. Johnson, and J.F. Marko, Nucleic Acids Res., 38, 6176 (2010).

[36] M.D. Stone, Z. Bryant, N.J. Crisona, S.B. Smith, A. Vologodskii, C. Bustamante, and N.R. Cozzarelli, Proc. Natl. Acad. Sci. U.S.A., 100, 8654 (2003).

[37] K. Yogo, T. Ogawa, M. Hayashi, Y. Harada, T. Nishizaka, and K. Kinosita, Plos One, 7 (2012).

[38] K. Terekhova, J.F. Marko, and A. Mondragon, Nucleic Acids Res., 42, 11657 (2014).

[39] M.X. Sun, T. Nishino, and J.F. Marko, Nucleic Acids Res., 41, 6149 (2013).

