## Supplemental Materials for "Measurement of the torque in braided DNAs using a thermodynamic Maxwell relation"

### using a thermodynamic Maxwell relation

Ke Ding[2], and John F. Marko[5,6]

**Supplementary Text to Figure 1:**

The extension shortening between Ca=0 and Ca=0.5 is used to calculate the intertether distance by the Pythagorean theorem. For Figure 1a, b (also listed below),

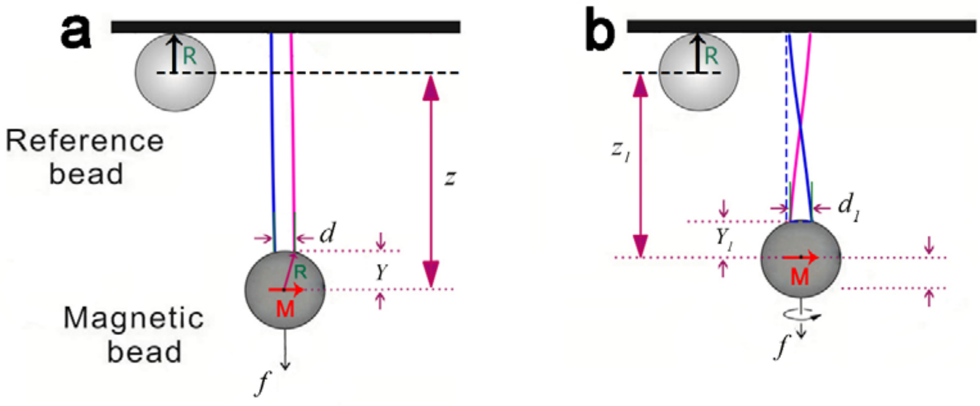


we use: $(z+R-Y_1)^2-(z_1+R-Y_1)^2=d^2$ $\quad$ $R^2-Y_1^2=\dfrac{d^2}{4}$

where $R$ is the radius of the magnetic bead, $z$ is the extension at Ca=0, and $z_1$ is the extension at Ca=1. The calculated distance $d$ in Figure 1f-h were 0.52, 0.62, and 1.06 µm, respectively.

#### Supplementary Method

**Theoretical calculations using the semi-flexible polymer model**

We compare our experimental data with the results obtained from a recently developed free energy model for a braid considering double helical DNAs as electrically charged semiflexible polymer [1,2]. For two braiding double helical DNAs under a constant holding force $f\hat{\mathbf{z}}$, the extension of the unbuckled part of the braid, which is directly coupled to the external force is obtained from the negative force derivative of the free energy. The net end-to-end extension of the braid can be obtained as a sum of contributions from the helically braided part ($z_b$) and the triangular end regions ($z_e$) (Note, the plectonemic region is decoupled from the external force and does not contribute to extension): $z = z_b + z_e$, where $z_b$ and $z_e$ are given as follows:

$$z_b = L_b cos\delta \left[ 1 - \frac{3}{4\sqrt{2}} \sqrt{\frac{k_B T}{fAcos\delta}} - \frac{k_B T \eta^{1/4}}{fAcos\delta} \left( \left( \frac{4k_B T\sqrt{\eta}}{fAcos\delta} \right)^2 - 1 \right)^{-1/2} sin\left( \frac{1}{2} tan^{-1} \sqrt{\left( \frac{4k_B T\sqrt{\eta}}{fAcos\delta} \right)^2 - 1} \right) \right] \quad \text{S1}$$

$$z_e = L_e cos\phi \left[ 1 - \sqrt{\frac{k_B T}{fAcos\phi}} \right] \quad \text{S2}$$

Here $L_b$ are $L_e$ are respectively the contour lengths of the helically braided part and the end regions, such that the total length of the DNA is divided into three parts: $L=L_b+L_e+L_p$, where $L_p$ is the length of DNA that forms braid plectonemes. The braid angle is given by $\delta$ whereas the end-angle, joining the braid with the end regions is given by $\phi$. The electrostatic contribution that drives an increase in the braid radius due to repulsion between the braiding strands is considered via $\eta$, which is the modulus of braid radii deformation given by the electrostatic potential. All these parameters that determine the mean field structure of the braid, at a given catenation and external force, are determined via a free energy minimization [1].

In Eq. S1, showing the extension of the helically braided part, the first term corresponds to the mean-field elastic extension, whereas the subsequent terms are fluctuation corrections. Transverse fluctuation of the braiding strands about their mean-field shape leads to a lower end-to-end extension, where fluctuations are suppressed by a higher external force and a stronger electrostatic modulus. Out of the four normal modes of fluctuations in a braid, three are dominated by the external force (the second term in Eq. S1), whereas one is controlled by the electrostatic modulus (third term in Eq. S1).

Similar to the case of the end regions, entropic elasticity of DNA leading to transverse fluctuations decrease the extension from its elastic straight-rod limit denoted in the first term of Eq. S2. The second term depicts the transverse fluctuations that are suppressed by the external force [1]. The extensions calculated from the numerical derivatives of the free energy is typically shorter by less than ~5% compared to Eqs. S1 and S2. This is because, although the mean-field plectoneme state is decoupled from force and does not contribute to end-to-end extension, the fluctuation corrections in the plectoneme structure are controlled by the external tension.

[1] Brahmachari,S. and Marko,J.F. (2017) Torque and buckling in stretched intertwined double-helix DNAs. *Phys. Rev. E* **95**, 052401.

[2] Brahmachari,S., Gunn,K.H., Giuntoli,R.D., Mondragon,A. and Marko,J.F. (2017) Nucleation of Multiple Buckled Structures in Intertwined DNA Double Helices. *Phys. Rev. Lett.* **119**, 188103.
